# Robust optical autofocus system utilizing neural networks trained for extended range and time-course and automated multiwell plate imaging including single molecule localization microscopy

**DOI:** 10.1101/2021.03.05.431171

**Authors:** J. Lightley, F. Görlitz, S. Kumar, R. Kalita, A. Kolbeinsson, E. Garcia, Y Alexandrov, V. Bousgouni, R. Wysoczanski, P. Barnes, L. Donnelly, C. Bakal, C. Dunsby, M.A.A. Neil, S. Flaxman, P.M.W. French

## Abstract

We present a robust, long-range optical autofocus system for microscopy utilizing machine learning. This can be useful for experiments with long image data acquisition times that may be impacted by defocusing resulting from drift of components, e.g. due to changes in temperature or mechanical drift. It is also useful for automated slide scanning or multiwell plate imaging where the sample(s) to be imaged may not be in the same horizontal plane throughout the image data acquisition. To address the impact of (thermal or mechanical) fluctuations over time in the optical autofocus system itself, we utilise a convolutional neural network (CNN) that is trained over multiple days to account for such fluctuations. To address the trade-off between axial precision and range of the autofocus, we implement orthogonal optical readouts with separate CNN training data, thereby achieving an accuracy well within the 600 nm depth of field of our 1.3 numerical aperture objective lens over a defocus range of up to approximately +/− 100 μm. We characterise the performance of this autofocus system and demonstrate its application to automated multiwell plate single molecule localisation microscopy.

## 1. Introduction

Optical microscopes using high numerical aperture objective lenses typically have a depth of field of the order of 1 μm. Thermal fluctuations and other causes can lead to mechanical drift of microscope components over time such that the region of interest (ROI) being imaged in the sample is not maintained within the imaging depth of field. This will compromise many experiments, including time-lapse imaging or super-resolved microscopy. In automated multiwell plate imaging and slide-scanning microscopes, this issue can be compounded with the sample holder deviating from flatness or not being perpendicular to the optical axis such that the sample goes out of focus when the instrument moves to imaging a different field of view. Thus, it is desirable to utilise an active form of focus correction (often called “autofocus”) where the sample or lens is moved axially to restore or maintain the sample ROI within the depth of field (DOF) of the objective lens. For automated multiwell plate microscopy techniques that require relatively long image data acquisition times, such as single molecule localisation microscopy (SMLM)^1,2^, an autofocus system is necessary to address both drift during the long image acquisition times when imaging over a multiwell plate and the well-to-well variation in distance of the coverslip from the objective lens.

There are two broad approaches to realize autofocus functionality. Image-based autofocus techniques, e.g. ^3^, analyse the images inherently acquired in the microscope, which can be brightfield or fluorescence images, and calculate image-based metrics that indicate the degree of defocus - and therefore the required movement of the objective lens or sample stage to bring the sample ROI back within the imaging DOF. The alternative approach, described as hardware-based autofocus or optical autofocus, entails the incorporation of a separate optical system in the microscope that measures how far the microscope coverslip/sample interface, e.g.^4^, (or other feature in the sample, which may include fiduciary markers^5^ such as gold beads) is from the objective lens and therefore provides a signal to control the movement of the objective lens or sample stage to bring the imaging system back to focus. These optical autofocus systems are typically limited in precision by diffraction. For extreme super-resolved microscopy experiments where resolutions approaching 10 nm are desired, a more precise axial localisation and drift control can be achieved using intensity and phase imaging of fiduciary nanoparticles with a wavefront-sensing device to determine defocus^6^.

While image-based autofocus systems do not require any modification of the microscope hardware, they can be slow, since they typically require the acquisition of multiple images. This can compromise biological samples since the additional images increase the light dose and therefore the possibility of photobleaching and/or phototoxicity. Moreover, image-based autofocus systems tend to work over a relatively narrow range close to the depth of field, which can be inadequate for automated multiwell plate imaging where axial variation in the distance to the well plate can be as much as ~250 μm across the plate. Supplementary Figure S1 shows a map of this axial variation measured for a typical 96 well plate (Brooks life Science Systems, MGB096-1-2-LG-L).

Hardware-based autofocus techniques can utilise a range of different principles to measure the sample-objective lens distance including confocality (e.g. measuring intensity of light reflected back from coverslip and focused though a pinhole^4^), or the measurement of the change in an image metric (such as displacement of images^7^ or centroid^8^) of an optical signal from the sample. The latter may be fluorescence emission^9^ or a (laser) beam that is focused onto and back reflected from the microscope coverslip^6^, or scattered from fiduciary markers^5^ (for which infrared radiation can be used to avoid cross-talk with visible fluorescent probes and to minimise phototoxicity). Techniques based on measuring defocus through measurements of the back-reflected beam size have an operational range that scales with the depth of field of the focused autofocus beam. This results in an inherent compromise between achieving high precision (which requires short depth of focus comparable to the imaging depth of field) or longer range (which results in lower precision). Furthermore, to distinguish whether the defocus is positive or negative, they may be set to measure distance from an offset position^10^ rather than when the autofocus laser beam is at focus, and this further reduces the range of the autofocus.

Alternatively, a readout of defocus can be realized by displacing the laser beam from the optical axis such that there is a change and/or shift of the intensity distribution of a back-reflected beam at the detector, which can be a camera or a position-sensitive detector (PSD, usually based on an array of photodiodes). Calibration of the variation of the position-based measurement with defocus can yield the defocus magnitude and sign.

The need for calculations of image-based metrics to quantify the defocus can be avoided by utilizing machine learning techniques to quantify the defocus from the signals recorded at the detector. Convolutional neural networks (CNN) have been trained to utilise speckle patterns^11^ recorded on a camera with the training sets being z-stacks acquired at known axial positions. Here we present a different approach using machine learning with CNN applied to analyse images of a back-reflected laser beam that change with the degree of defocus of the microscope.

## 2. CNN-based optical autofocus

Here we report the development of a hardware-based optical autofocus system that we have developed for automated multiwell plate super-resolving microscopy based on *easySTORM^12^*, which is a cost-effective implementation of direct stochastically switched optical reconstruction microscopy (dSTORM)^13^ that utilizes low-cost multimode diode lasers and provides large (> 100 μm) fields of view and can be adapted to install on almost any epifluorescence microscope. The image data acquisition for *easySTORM* (and other single molecule localisation microscopy (SMLM) techniques) typically requires several minutes (e.g. 150 seconds to acquire 5000 frames for a single STORM image), which is long enough for the system to be impacted by thermal drift if the local temperature is not well-controlled. In addition, multiwell plate imaging of sample arrays typically requires refocusing at each new well. Accordingly, we implemented the set-up depicted in Figure 1, which essentially images the intensity distribution of an infrared beam (from a diode laser emitting at 830 nm) that is focused onto the microscope coverslip and imaged back to the autofocus camera, positioned in a conjugate plane to the microscope coverslip.

**Figure 1.**
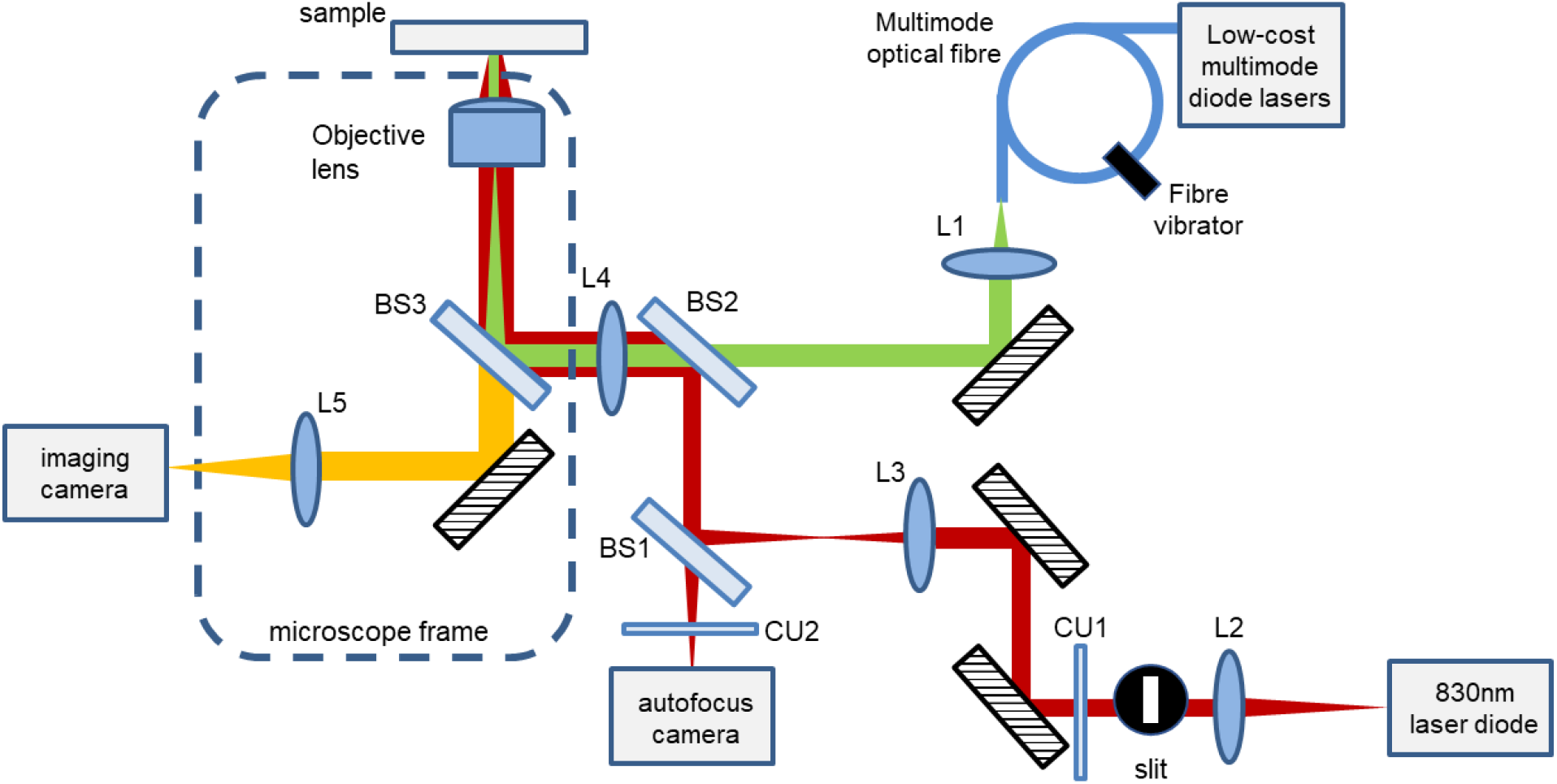
Schematic of easySTORM microscope utilizing optical autofocus. The objective lens was an oil immersion 1.3 NA 100x lens with a depth of field of 600 nm. The focal lengths of the other lenses were L1: 50mm, L2: 18.4mm, L3: 100mm, L4: 200mm, L5: 180mm. BS are beam splitters and CU are clean-up filters. For further experimental details, please see the Supplementary Information.

Supplementary Figure S2 shows an exemplar z-stack of the autofocus camera images acquired as the microscope coverslip is translated through the focus of the microscope objective lens. The extent of defocus can be determined analytically by calculating image metrics (such as peak image intensity or standard deviation) of the images acquired at the autofocus camera that will typically change systematically as the coverslip is moved from focus. By acquiring a z-stack of images at the autofocus camera for known values of defocus (i.e. distance of the coverslip from the focal plane), a look-up table (e.g. for specific image metrics) can be devised to enable the degree of defocus to be determined from a single autofocus camera image. Alternatively, we can use a number of such experimentally acquired through-focus z-stacks to train a CNN model that can be used to determine the degree of defocus from an autofocus camera image.

For both deterministic and machine-learning approaches, the precision of localising the focus position depends on the confocal parameter of the focused infrared beam, which in turn depends on the focal length of the objective lens and the diameter of the incident beam. Usually this results in a trade-off between precision and range for such autofocus systems, since the defocused image rapidly expands beyond the camera sensor area as defocus is increased. We have mitigated this compromise by adding a slit after the lens (L2) that collimates the near infrared diode laser, which has the effect of introducing two simultaneous confocal parameters due to the different beam diameters in orthogonal directions. Using a slit with an aspect ratio of 3:1 and a two-step correction process, this provides us with an autofocus precision of less than 200 nm (benefiting from the short-range confocal parameter measurements) over a range of +/− 100 μm (enabled by the by the long-range confocal parameter measurements). An exemplar autofocus camera z-stack is shown in Supplementary Figure 2. In our experience, this range of +/−100 μm is sufficient for multiwell plate imaging, noting that when following a snake-like path through adjacent wells, the well-to-well variation is usually much less than 100 μm.

We found that if an image metric is used to deterministically calculate the defocus from the images acquired at the autofocus camera, it typically requires calibration before each experiment, since the precise images recorded by the autofocus camera can themselves be impacted by small changes in alignment, e.g. resulting from thermal drift, which can occur during an extended experiment. We therefore investigated whether the CNN-based readout would be more robust. In a manner like that of reference [5], a CNN model was trained on z-stacks of images acquired by the autofocus camera at known defocus. Figure 2 depicts the processes for acquiring the training data and applying the CNN to determine the defocus.

**Figure 2.**
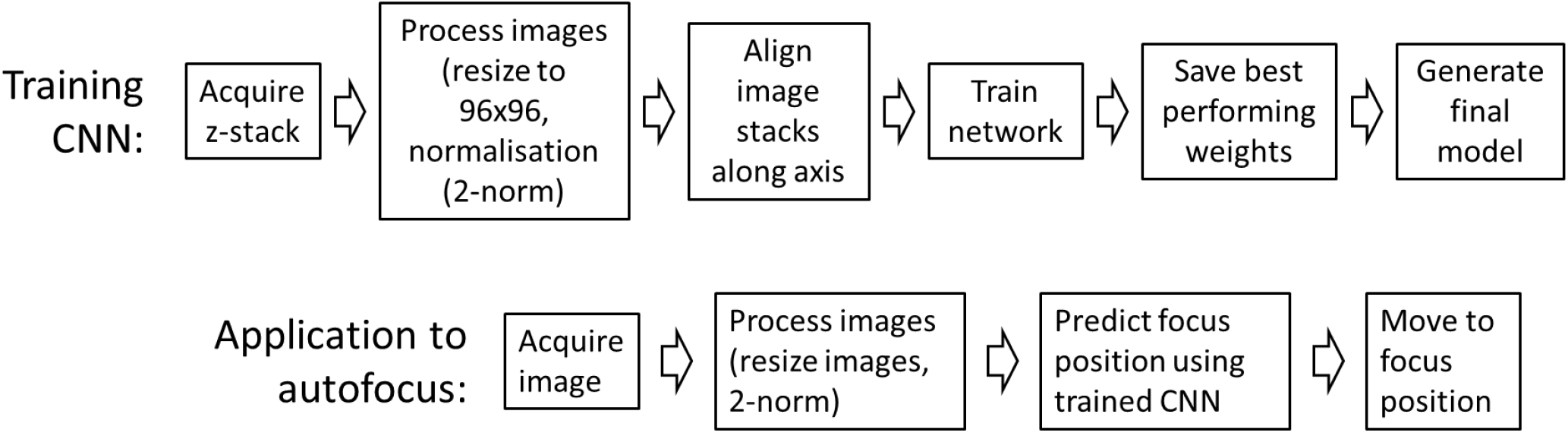
Schematic of processes to train CNN for optical autofocus

For the CNN we have utilized the MobileNetV2 architecture^14^ as the base structure for the model, which is implemented as part of the TensorFlow^15^ 2.1 package installation in Python. The top of the model was not included and was replaced with a global average pooling 2D layer followed by a dense layer with a scaler output. This architecture allows for a regression prediction of the defocus value. The model has an inverted residual architecture with 157 layers in total and makes use of Depthwise Separable Convolutions and Linear Bottlenecks^13^ to reduce the memory and operation requirement during training. The outputs of these layers were then input into a max pooling layer, whose output was converted into a 1-dimensional array using a flatten layer before being passed to a series of densely connected layers to give a scalar output.

Each CNN training data set comprised 100 z-stacks of images saved as OME TIF stacks for which the corresponding z-stage position was recorded in the metadata for each image. As discussed below, the z-stacks were acquired with 1 μm axial sampling over +/−100 μm or with 50 nm sampling over +/−10 μm. This image data acquisition takes about 3 hours. For the CNN training, each autofocus camera image in each of the z-stacks was rescaled and binned using the OpenCV package in Python into a NumPy array of shape (96×96×1) to be used as the input shape for the model. This input data was normalised using the Euclidean norm. The model was trained using an MSI GeForce RTX 2080 SUPER VENTUS XS OC GPU.

In order to determine the “ground-truth” defocus offset for each image within a data stack, the stacks had to be aligned prior to training with respect to the focal plane, defined as the zero-defocus position. To determine the zero-defocus position for each z-stack, the autofocus camera images were averaged along the pixel rows to provide a one-dimensional vertical intensity projection for each z-value and a Savitzky-Golay filter was passed along these averaged intensity projections. The standard deviation of the squared absolute value of the Fourier transform (power spectrum) of these smoothed profiles was then plotted as a function of z to yield a smoothly varying function of z with a minimum at the zero-defocus position. An example of this function is presented in Supplementary Figure S3. All the images in the z-stack were then labelled according to their distance from the zero-defocus image.

The defocus labelling for each image was then normalised (i.e. centred on the mean and divided by half the total range) for training the CNN. This was repeated for all the data stacks used for training and 10% of the data stacks were removed from each day’s training data to be used as validation datasets. The optimization algorithm Adam^16^ was used for training with a learning rate of 0.0001 and the mean squared error was used as the loss function. The loss was defined in the normalised space. The validation loss was recorded after each epoch and the weights of the model were only saved after each epoch if the performance of the model improved on the previous best performing epoch. Thus, only the best performing model’s weights for the validation were saved and used after training. This technique was used as a form of early stopping using checkpoints to prevent the model from overfitting to the training data.

After training the CNN, which takes approximately 3 hours, the application of the autofocus requires the acquisition of a single autofocus camera image, from which the CNN can determine the distance of the coverslip from the focal plane of the objective lens. Following a prediction of the defocus in normalised units, this is rescaled to the actual defocus offset (i.e. in μm) and then the z-stage can be directed to move the coverslip back to focus - or to a pre-programmed offset if imaging at a plane away from the coverslip is desired.

We observed that the precision to which the CNN can localize the focal position is limited by the axial sampling of the training set, which should therefore be finer than the required axial resolution. However, since the data volumes and CNN training times increase with the sampling density, over-sampling should be avoided. In the direction parallel to the long side of the slit (corresponding a “short-range confocal parameter”) we found that the recorded image was useful up to a range of ~+/− 10 μm, beyond which the beam overfills the camera sensor and the signal to noise ratio is no longer sufficient. Since the depth of field of the fluorescence imaging is ~0.6 μm, we acquired training data at 50 nm axial sampling over this 20 μm range. The readout interval of the axial microscope stage was 10 nm. If we were to continue sampling at 50 nm over the full 200 μm range, this would greatly increase the training data volume and computation time with no benefit. The autofocus long confocal parameter (in the orthogonal plane) was ~9 times larger and for this we acquired a second CNN “long-range” training set with 1 μm axial sampling over +/− 100 μm.

Figure 3 illustrates the performance of this CNN-controlled autofocus, which was studied by recording 10 z-stacks of images at different axial displacements on the autofocus camera. The top row of Figure 3(a, b, c) illustrate the performance of the short-range CNN model and the bottom row of Figure 3(d, e, f) illustrates the performance of the long-range CNN model. For each z-stack, the “true” axial location of the focal plane was determined by calculating the standard deviation of the power spectrum distribution of the autofocus camera intensities averaged across the pixel rows (corresponding to the short-range autofocus axis) for each image along this z-stack, in order to identify the z-value at which it was minimized. Then the value of defocus predicted from each autofocus camera image in the z-stack using the CNN is plotted against the “true” defocus value for the data corresponding to the short-range (a) and long-range (d) data. Figures 3(b, e) show the mean of the residual errors (i.e. CNN predicted value – “true” defocus value) for the 10 z-stacks as a function of z. The error in the mean residual is the standard deviation of the 10 residual plots at each data point. Figures 3(c, f) show the plots of the training (black) and validation (red) loss of the CNN for the short-(c) and long-(f) range training data as a function of epoch. Thus Figure 3 shows that better than ~100 nm accuracy is typically achieved over the +/−10 μm range when the short-range training data set is used, with a precision of <100nm and that <1 μm accuracy is achieved over most of the 200 μm range of the long-range training data set with a precision of <1 μm.

**Figure 3.**
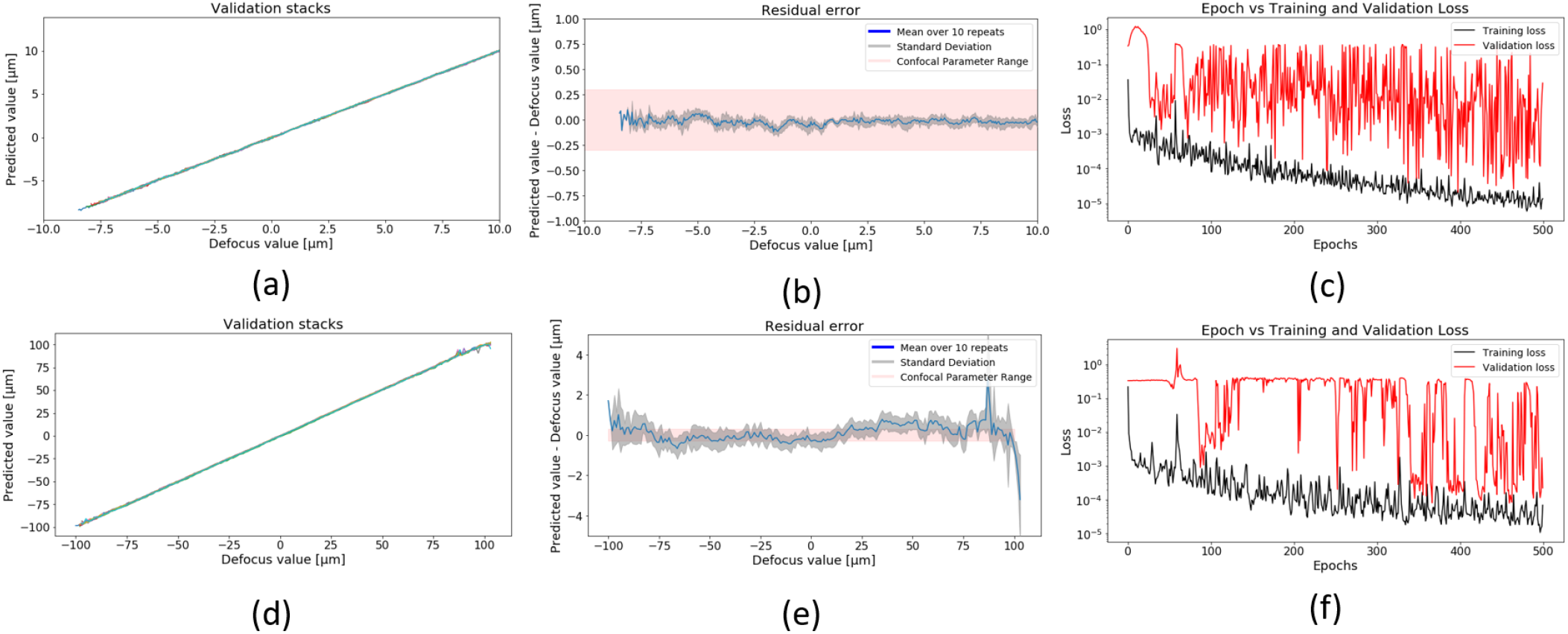
Plots of predicted vs actual defocus for 10 through-focus z-stacks, with each z stack plotted in a different colour (a, d), with the corresponding mean of the residual errors (b, e) resulting when using the short-range and long-range CNN training sets acquired and validated on data on the same day. Figures (c, f) show plots of the changing training and validation loss with increasing epochs during training for the short range and long-range CNN training sets respectively.

By combining these CNN training sets as illustrated in Figure 4, we can use the resulting aggregate CNN training set to provide the long-range autofocus capability but achieve the short range precision by applying the CNN defocus determination twice: the first application of the long range CNN training set brings the system to within the range (i.e. <10 μm from focus) where the short-range training set is valid and this can then determine the focus to <200 nm.

**Figure 4.**
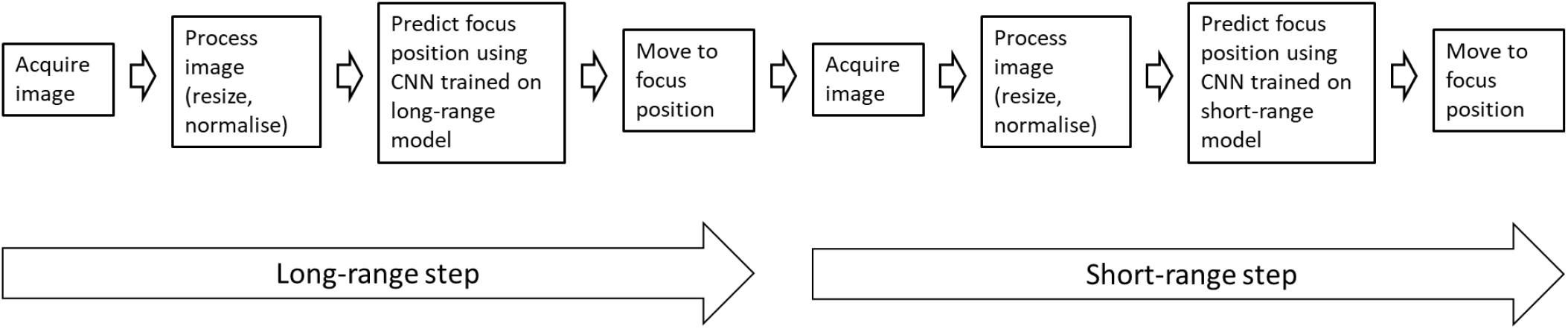
Diagram of data process stages for two-step autofocus

## 3. Impact and mitigation of thermal/misalignment system drift

Unfortunately, the thermal variations and other factors that can lead to defocusing of the microscope can also lead to errors in optical autofocus systems themselves. For commercial microscopes designed to incorporate an optical autofocus, the manufacturers typically engineer a compact optical autofocus system incorporating a position-sensitive detector or camera close to the microscope objective lens, so that the relatively short optical paths are less impacted by this “optical system drift”. For microscopes to which an external autofocus system is added, as is the case here, this system drift remains an issue as illustrated in Figure 5, which shows the errors in the autofocus that can occur when the CNN training data has been acquired even just one day earlier than its application. Here we measured the performance of the autofocus system as described for Figure 3, but we applied the CNN using training data measured on one day to defocus measurement data acquired on subsequent days.

**Figure 5.**
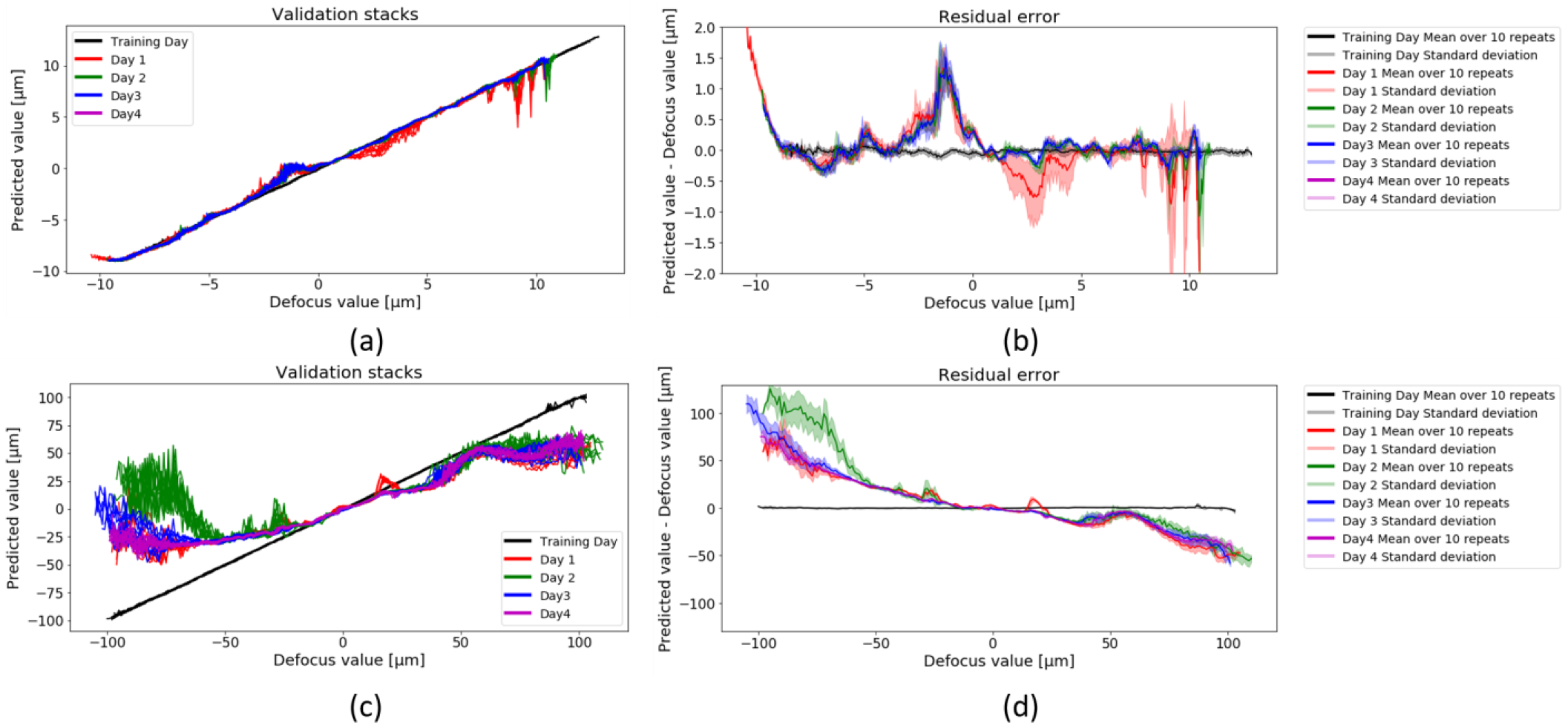
Plot of predicted vs actual defocus (a, c) and the corresponding mean of the residual errors (b, d) when using the short-range and long-range CNN training sets acquired (separately and sequentially) on one day for measurements undertaken on different subsequent days. Figures (a) and (c) show data from multiple days with 10 repeats for each day, with all data from a single day plotted in the same colour.

Since the acquisition of training data and the training of the CNN takes several hours, it is not practical to train the CNN with “fresh” data every day. Furthermore, for experiments lasting many hours, such as automated multiwell plate SMLM, this optical system drift can occur during a single experiment and degrade the experiment even using a freshly trained CNN. We reasoned that we could mitigate the impact of the optical system drift on the CNN autofocus system by expanding the training data to include such system variation. We have found empirically that using an aggregate CNN training data combining autofocus camera image z-stacks acquired on 6 separate days over the course of a 10 day period was sufficient to provide robust prediction of the defocus (i.e. less than the 600 nm imaging depth of field) on a subsequent day, as illustrated in Figure 6.

**Figure 6.**
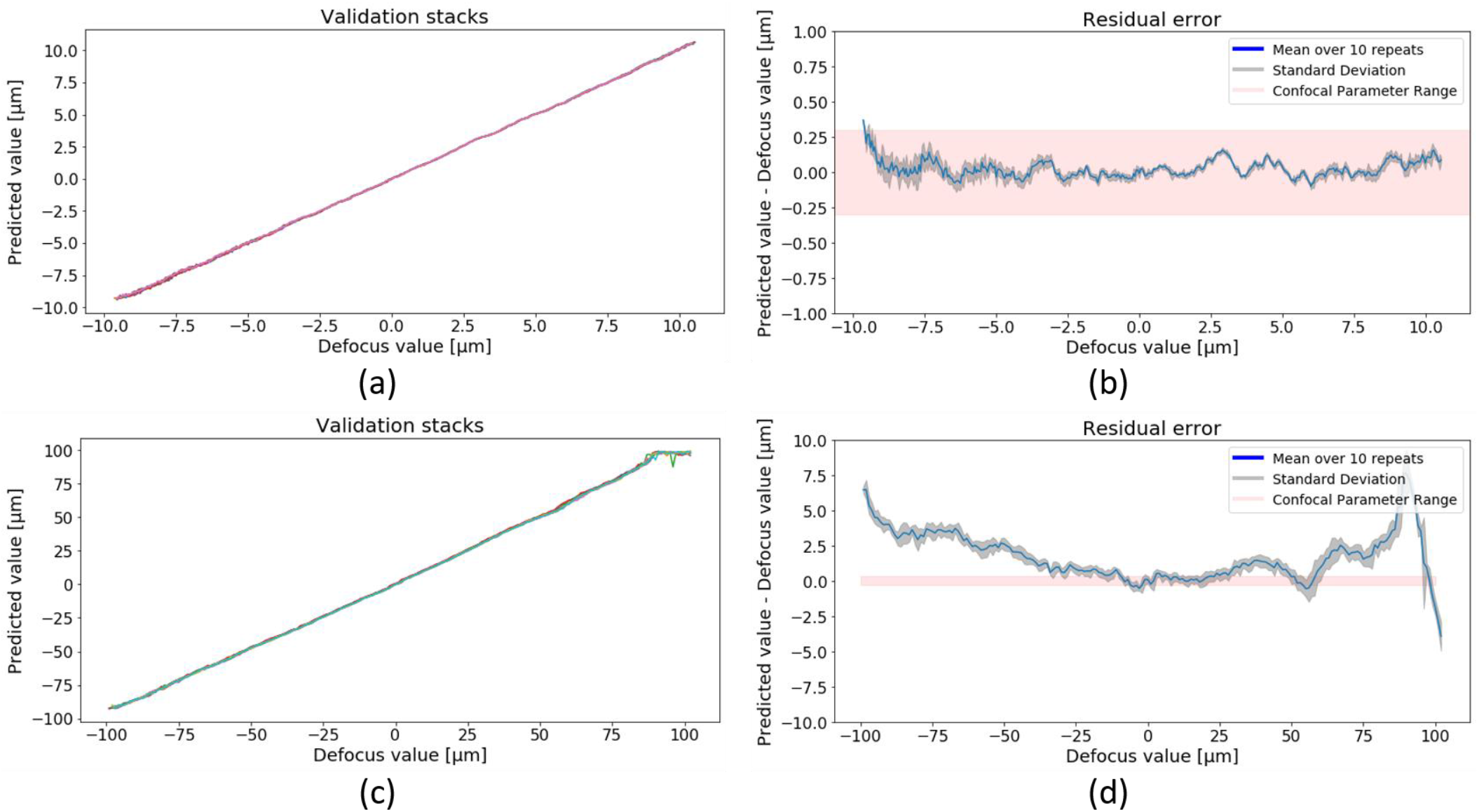
Plot of predicted vs actual defocus for 10 through focus z-stacks, with each z stack plotted in a different colour (a, d), and the corresponding mean of the residual errors (b, d) when using aggregate short-range and long-range CNN training sets that incorporate training data sets acquired over a period of 10 different days for measurements undertaken on a subsequent day.

To further quantify the performance of this CNN-based autofocus system, we undertook a systematic study for which we measured the error in determining the focal position starting from a range of different defocus positions using the aggregate CNN trained over 10 previous days. For each defocus starting point, we repeated the measurement 10 times and Figure 7 shows the resulting histograms of the “autofocus error” when using the short-range and long-range aggregated CNN training sets as well as the performance achieved using the two-step autofocus. The short range autofocus reproducibly brings the coverslip back to the focal plane back within the (~600 nm) depth of field of the fluorescence microscope over an axial range of +/− 10 μm as expected while the long range autofocus brings the coverslip back to within ~2 μm of focus over an axial range of +/− 100 μm. When combining the long- and short-range focus in the two-step approach, Figure 7(c) shows that the coverslip is returned to within +/−200 nm of focus (well within the fluorescence depth of field) for a defocus range over +/− 100 μm. This CNN-based autofocus trained over multiple days has sufficient range and precision for multiwell plate imaging applications, including easySTORM HCA.

**Figure 7.**
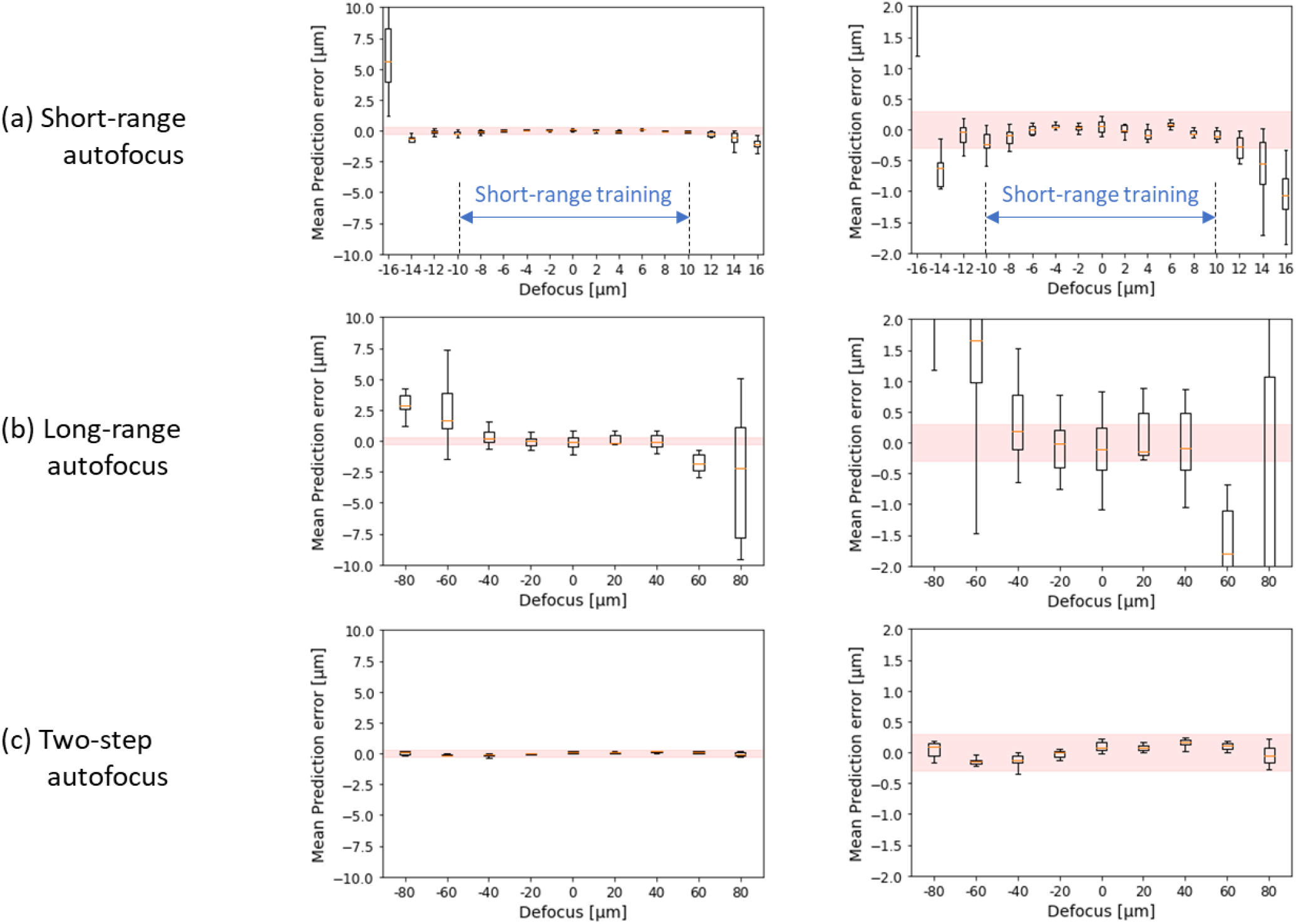
Plots of prediction error vs initial defocus for 10 measurements at each defocus position sequentially applying the short-range and long-range CNN training sets acquired over 10 previous days. The red band corresponds to the ~600 nm depth of field of the fluorescence microscope. The right-hand column shows same data as left-hand column on expanded vertical scale

## 4. Application of CNN-based optical autofocus system to multiwell plate imaging

To evaluate the performance of this autofocus approach for multiwell plate imaging, we developed a methodology that can be applied to any automated multiwell plate microscope. This entails automatically imaging a multiwell plate of sub-resolution fluorescent bead samples arrayed in each well and, using the PSFj^17^ software published previously that automatically calculates the distance from focus of sub-resolution beads from acquired z-stack images, we calculated the mean displacement of the beads in each well from the focal plane of the objective lens. Figures S4 and S5 show the results obtained using a 96 well plate (Brooks life Science Systems, MGB096-1-2-LG-L) arrayed with 100 nm diameter fluorescent beads (TetraSpeck, T7279). These results are displayed as maps of this mean displacement for each well and a histogram of these values, acquired for a horizontal snake acquisition track (Figure S4) and a vertical snake acquisition track (Figure S5). It is apparent in figure S4 that the multiwell plate is tilted along its long dimension such that the short range autofocus fails after imaging one well along the horizontal snake path because the displacement is more than the 10 μm range of the training data. The long range autofocus acquires image z-stacks in all of the wells and the standard deviation of the histogram of these mean displacement values is 1.17 μm when acquired in a horizontal snake pattern. When the two-step autofocus is used in a horizontal snake pattern, z-stacks of bead images are acquired in all wells and the standard deviation of the histogram of these mean displacement values is 0.21 μm (Figure S4). Figure S5 shows that, when acquiring the image z-stacks following a vertical snake path, the system acquires useable images in most of the wells with the short range autofocus and with the long range autofocus. The two step autofocus acquires image z-stacks all the wells with a standard deviation of the histogram of these mean displacement values of 0.16 μm, which is superior to the values of 0.26 μm for the short range and 0.98 μm for the long range autofocus systems respectively for the vertical snake. We note that this methodology to quantify the performance of an autofocus system for multiwell plate imaging does not rely on any properties of our autofocus system and can be generally applied to evaluate the performance of the autofocus of any automated microscopy system.

To illustrate the application of our two-step CNN-based autofocus to automated multiwell plate SMLM, we applied it to automated easySTORM of fixed cells arrayed in a 96 well plate, for which the details of the sample preparation can be found in the Supplementary information.

Figure 8 shows an exemplar data set of automatically acquired wide-field epifluorescence and easySTORM images acquired on an Olympus IX81 microscope with a motorised x-y stage (Marzhauser SCAN IM 112×74, controlled using the Marzhauser TANGO controller) incorporating the autofocus system represented in Figure 1. In this experiment WM2664 melanoma cells arrayed in 6 × 4 wells were either mock treated (control, bottom row), or incubated with siRNA targeting the ARHGEF9 (top rows) and Anillin proteins. The cells were plated on fibronectin for 2 hours before fixation, and then stained with phalloidin to label F-actin, and anti-Paxillin antibody to label the focal adhesions (FA). The acquired image data were processed using ThunderSTORM implemented on our high-performance computing (HPC) cluster^18^.

**Figure 8.**
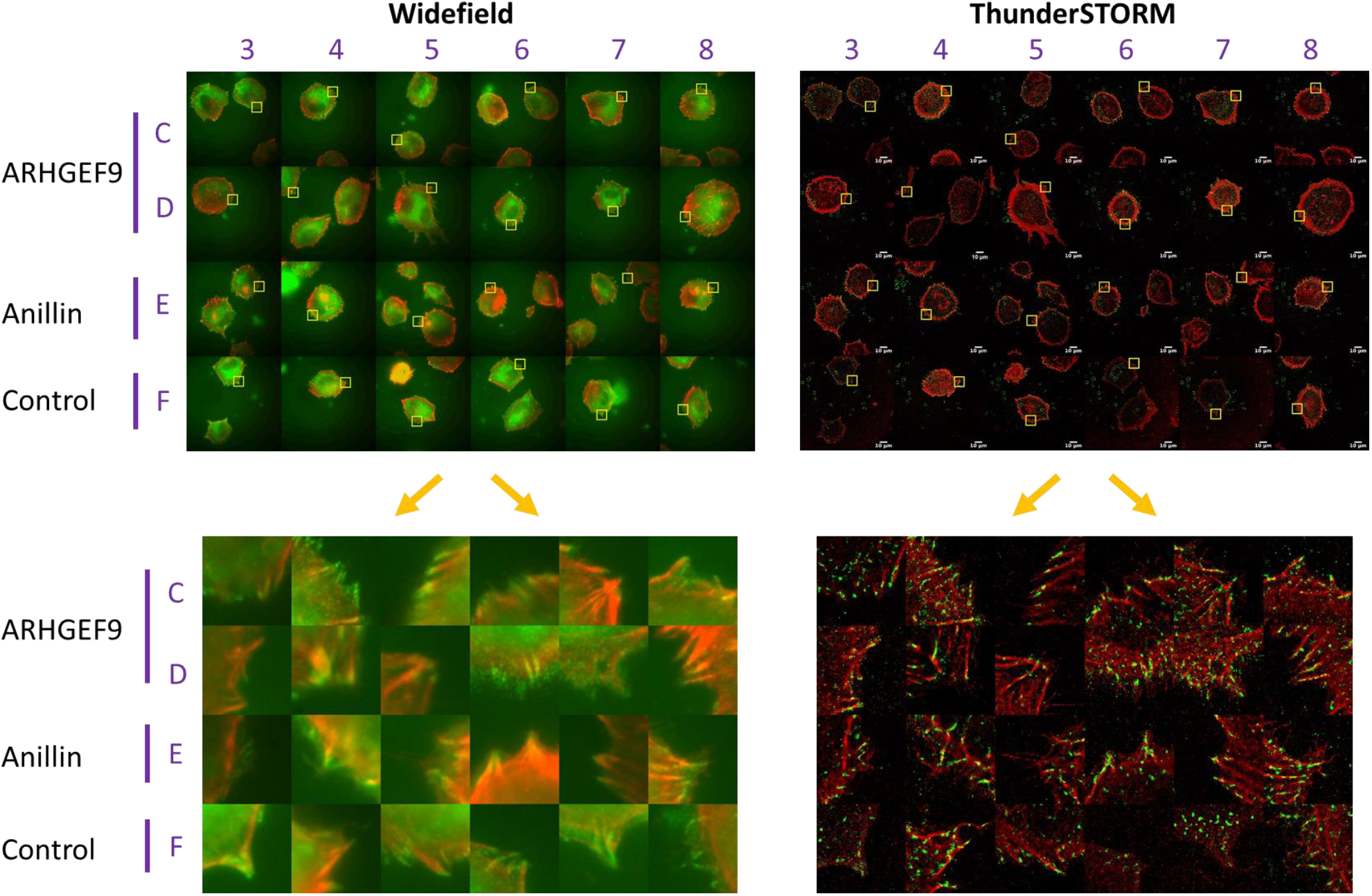
shows automatically acquired wide-field epifluorescence and easySTORM images of an array of WM2664 melanoma cells arrayed in 6×4 wells of a 96-wel plate that were either mock treated (control, bottom row), or incubated with siRNA to knock down the ARHGEF9 (top two rows) and Anillin (third row) proteins. The top figures show thumbnail images of a typical field of view (125.8×125.8 μm) in each well and the bottom figures show expanded images of the indicated (yellow boxes) regions of interest (12.8×12.8 μm).

The wide-field images show some gross differences in cell morphology (upper left), and differences in the distribution of F-actin and FAs (expanded regions of interest, bottom left) are apparent. The *easySTORM* images on the right reveal that in control cell small, nascent FAs have formed at the cell periphery as indicated by the formation of punctate Paxillin foci (green). The levels of polymerized F-actin are low around these structures and the organization of Paxillin and actin is characteristic of nascent FA structures undergoing turnover in dynamic protrusions with high levels of actomyosin flow. In contrast, in both ARHGEF9 and Anillin depleted cells, Paxillin is organized into elongated structures that align along long filaments of F-actin. Remarkably, multiple Paxillin-positive structures are found aligned in a single “row” along single actin fibers. These data demonstrate that the autofocus system can enable automated multiwell plate easySTORM microscopy to provide super-resolved images

Figure 9 shows a second proof-of-principle demonstration of *easySTORM* HCA, here applied to image the uptake of bacteria in human monocytic THP-1 cells (ATCC) derived from an acute monocytic leukaemia patient cultured in Roswell Park Memorial Institute-1640 (RPMI), Sigma-Aldrich). THP-1 cells are widely used to model macrophage/pathogen interactions^19^. A multiwell plate was arrayed with THP-1 cells that were incubated with heat-killed *Streptococcus pneumoniae* labelled with AF488 for up to 60 minutes in 15-minute intervals, before washing to remove non-bound bacteria. The cells were fixed and stained with a primary monoclonal antibody for alpha tubulin labelled with AF647. The reconstruction of the automatically acquired image data was performed using WindSTORM^20^ implemented on an HPC cluster. Exemplar automatically acquired easySTORM images are presented with expanded images of cells from selected wells. As the incubation time of the bacteria is increased, there is a subsequent increase in bacterial uptake into the cells. It can be observed that with the increased internalization, there is an increased overlap of bacteria with the tubulin filaments. Work is underway to quantify the degree of co-localization of the bacteria with the tubulin, which appears to correlate with incubation time.

**Figure 9.**
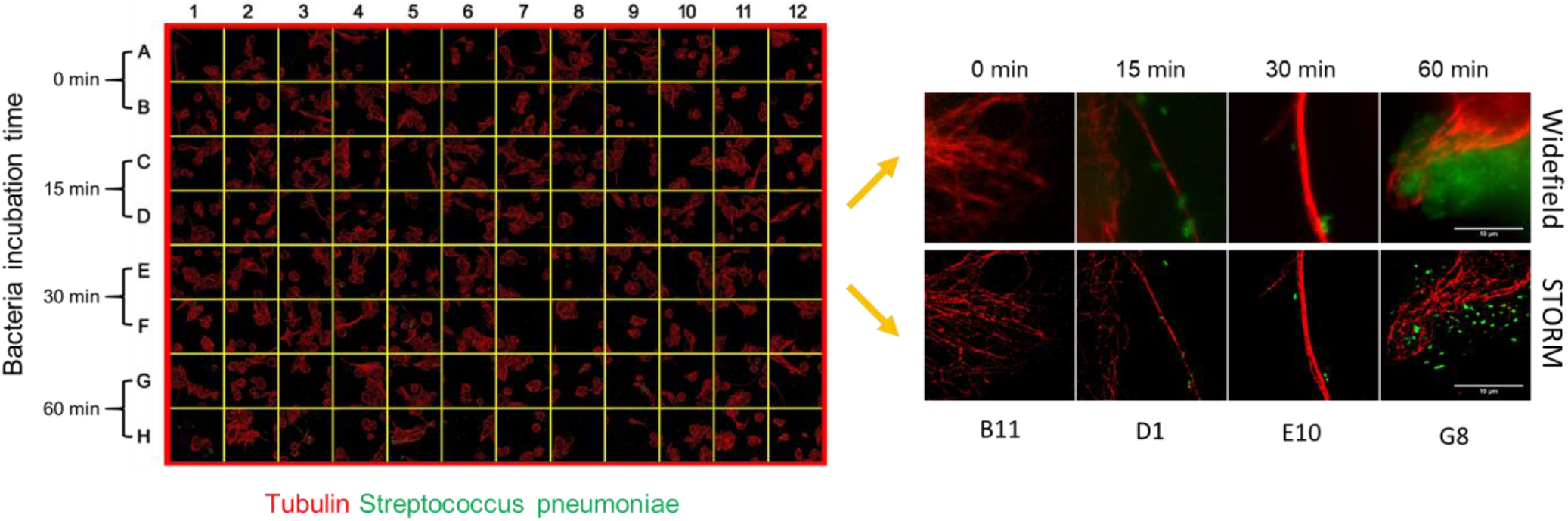
shows an array of automatically acquired easySTORM images of THP-1 cells incubated with Streptococcus pneumoniae bacteria and fixed for imaging. The alpha-tubulin in the THP-1 cells is labeled with AF647 (red) and the bacteria are labelled with AF488. An array of STORM images is shown in their respective wells (left) with exemplar expanded wide-field and easySTORM images (right) illustrating the change in bacteria distribution with increasing incubation time.

## Discussion

We have presented the design and performance of a hardware-based optical autofocus system that utilises a CNN-based readout to enable automated imaging. Once the CNN model is trained, the operation of the autofocus requires only 300 ms. This optical autofocus system was specifically designed for automated multiwell plate imaging using relatively time-consuming modalities such as SMLM for which it is necessary to refocus at each new field of view and to maintain focused imaging over several hours. We typically apply the optical autofocus every 200 frames to maintain focus during an SMLM acquisition.

We found that a CNN trained on data acquired on a single day could not be used successfully on subsequent days. To the best of our knowledge, although CNN-based machine learning has been applied to a range of optical instruments including a hardware-based autofocus system^11^, quantitative analysis of the impact of optical system drift over time has not been previously presented. The sensitivity of the CNN-based autofocus to this optical system drift, and the 6 hours required for acquisition of training data and training the CNN, makes it impractical to train a new CNN for every experiment. Furthermore, we observed that thermal drifts could even compromise its operation over the duration of a 2-hour multiwell plate imaging experiment undertaken on the same day as the CNN was trained.

We note that using an algorithmic approach based on calculating image metrics of the image z-stack acquired through focus could be used to generate a “fresh” look-up table to read out defocus from the autofocus camera images in a few minutes. However, this would not avoid the impact of optical system drift during a long experiment.

We observed that this optical system drift can be mitigated by training the CNN with data sets aggregating measurements over multiple days to “capture” this drift and incorporate it in the training model. In practice, CNN training data aggregated over 10 days provided robust autofocus over weeks of operation. Eventually, the optical system may undergo a significant perturbation not captured within the aggregate training data, e.g. due to the instrument being knocked or the system being reassembled. We found that the performance of the autofocus can be quickly regained by using “transfer learning”^21^ if the optical system is brought back into “proper alignment” such that the autofocus laser beam is aligned along the optical axis of the objective lens.

Transfer Learning involves using the transfer of knowledge from a previously trained model to make predictions on a new but similar task or dataset. The transfer of knowledge can reduce the amount of new training data that is required to achieve the same prediction accuracy as achieved previously for the new dataset. We found that transfer learning could restore the utility of the aggregate CNN model by incorporating a single additional training data set. For this, 10 z-stacks of autofocus camera images were acquired in a single day, of which 9 were used as a training dataset and one was used as a validation dataset. In order to transfer the learned weights from the previous model, the first 100 layers of the MobileNetV2 model based on the previous aggregated training data were copied and then frozen such that the model’s weights for these layers could not be changed. This allows the model to fine tune to the new system alignment without changing the generic features that have already been trained in the lower layers. The learning rate for the transfer learning was reduced to 0.00001 and the Adam optimiser algorithm was used again. The previously trained model was used as the base model and the model was trained for an additional 50 epochs. After training, the performance of the model was fully recovered to provide the same precision over the same range of defocus as before and the same performance over subsequent days.

This demonstrates the robustness of this CNN-based autofocus system, which has, to date, been used over a timescale of three months, during which time it was realigned twice. This optical autofocus system is now part of a “work-horse” automated easySTORM microscope that is being applied to a range of cell biology studies. The design can be implemented in the excitation light path of most epifluorescence microscopes provided that the main dichroic beamsplitter will reflect the (infra-red) autofocus laser beam.

## Conclusions

We have demonstrated a robust optical autofocus system that can be added to an existing optical microscope and here is introduced into the excitation light path of an epifluorescence microscope. We have addressed the trade-off between precision and range of this autofocus approach through the use of a slit to adjust the confocal parameter of the autofocus laser beam in orthogonal directions and the implementation of a two-step (long-range then short-range) determination of defocus. We have implemented a machine learning technique based on CNN to determine the displacement of the coverslip from the focal plane of the objective lens and shown that this two-step process works over a range of +/−100 μm with a precision well within the depth of field (~600 nm) of the objective lens.

We have illustrated, for the first time to our knowledge, how the performance of the CNN-based readout degrades over time as the precise configuration of the instrument changes the recorded optical autofocus signal and we have shown how this can be mitigated for future operation of the autofocus system by acquiring training data over multiple days (i.e. over the timescale on which these changes are occurring).

We have also presented a simple methodology to characterise the performance of any autofocus technique used with automated microscopes. To demonstrate that this autofocus system is practically useful, we have undertaken the automated acquisition of arrays of easySTORM images of biological samples, illustrating the potential to study the impact of specific siRNA gene knockdowns on the structure of focal adhesions in melanoma cancer cells and to study the incubation of bacteria in cells.

## Acknowledgements

Significant components of the instrument reported here were co-designed and fabricated by Simon Johnson and Martin Kehoe in the Optics instrumentation facility of the Physics Department at Imperial College. We gratefully acknowledge funding from the Development Fund of the Cancer Research UK ICR/Imperial Convergence Science Centre and Research England GCRF Institutional Award as well as the Imperial College London Impact Acceleration Accounts supported by the Biotechnology and Biological Sciences Research Council (BBSRC EP/R511547/1) and the Engineering and Physical Sciences Research Council (EPSRC EP/R511547/1). We also acknowledge funding from EPSRC (EP/V002910/1). SK is supported by funding from the Francis Crick Institute, which receives its core funding from Cancer Research UK (FC001003), the UK Medical Research Council (FC001003) and the Wellcome Trust (FC001003). CJB and VB are funded by a Cancer Research UK (CRUK) Programme Foundation Award and the Stand Up to Cancer campaign (C37275/A20146). JL acknowledges PhD studentships from EPSRC, RW acknowledges a PhD studentship from BBSRC. For the purpose of open access, the authors have applied a CC BY public copyright licence to ay Author Accepted manuscript version arising from this submission.

## Notes

### Competing Interest Statement

The authors have declared no competing interest.

## References

1 Holden, S. J., Pengo, T., Meibom, K. L., Fernandez Fernandez, C., Collier, J., & Manley, S. “High throughput 3D super-resolution microscopy reveals Caulobacter crescentus in vivo Z-ring organization,” Proc. Natl. Acad. Sci. U.S.A. 111, 4566–4571 (2014).

2 Beghin, A., Kechkar, A., Butler, C., Levet, F., Cabillic, M., Rossier, O., … Sibarita, J. B. “Localization-based super-resolution imaging meets high-content screening,”. Nature Methods, 14, 1184–1190 (2017).

3 Y. Sun, S. Duthaler, and B. J. Nelson, “Autofocusing in computer microscopy: Selecting the optimal focus algorithm,” Microscopy Research and Technique, vol. 65, pp. 139–149, (2004).

4 Y. Liron, Y. Paran, N. G. Zatorsky, B. Geiger, and Z. Kam, “Laser autofocusing system for high-resolution cell biological imaging,” J Microsc, vol. 221, no. 2, pp. 145–151, 2006.

5 Coehlo et al, “Ultraprecise single-molecule localization microscopy enables in situ distance measurements in intact cells”, Sci Adv 6 (2020) eaay8271

6 P. Bon, N. Bourg, S. Lécart et al.” Three-dimensional nanometre localization of nanoparticles to enhance super-resolution microscopy”. Nat Commun 6, 7764 (2015). https://doi.org/10.1038/ncomms8764

7 Liao et al, “Single-frame rapid autofocusing for brightfield and fluorescence whole slide imaging” Biomedical Optics Express 7(2016) 4763

8 Liu, Chien-Sheng, et al. “Novel fast laser-based auto-focusing microscope.” SENSORS, 2010 IEEE. IEEE, 2010.

9 Silvestri et al, bioRxiv http://dx.doi.org/10.1101/170555

10 Gu, Chao-Chen, et al. “A High Precision Laser-Based Autofocus Method Using Biased Image Plane for Microscopy.” Journal of Sensors 2018 (2018)

11 Henry Pinkard, Zachary Phillips, Arman Babakhani, Daniel A. Fletcher, and Laura Waller, “Deep learning for single-shot autofocus microscopy,” Optica 6, 794–797 (2019), https://doi.org/10.1364/OPTICA.6.000794

12 Kwakwa et al., “easySTORM: a robust, lower-cost approach to localisation and TIRF microscopy”, J. Biophotonics 9 (2016) 948–957, http://dx.doi.org/10.1002/jbio.201500324

13 Heilemann, M., van de Linde, S., Schüttpelz, M., Kasper, R., Seefeldt, B., Mukherjee, A., … Sauer, M. (2008). Subdiffraction-resolution fluorescence imaging with conventional fluorescent probes. Angewandte Chemie (International Ed. in English), 4733, 6172–6176.

14 Sandler, M.; Howard, A.; Zhu, M.; Zhmoginov, A.; Chen, L.C. MobileNetV2: Inverted Residuals and Linear Bottlenecks. In Proceedings of the IEEE Conference on Computer Vision and Pattern Recognition, Salt Lake City, UT, USA, 18–22 June 2018; pp. 4510–4520, arXiv:1801.04381

15 Abadi et al. “TensorFlow: Large-Scale Machine Learning on Heterogeneous Distributed Systems,” 14 March 2016, arXiv:1603.04467

16 Kingma and Ba, “Adam: A Method for Stochastic Optimization,” 22 december 2014, arXiv:1412.6980

17 P. Theer, C. Mongis and M. Knop, “PSFj: know your fluorescence microscope”, Nat Methods, 11 (2014) 982

18 Munro, I., Garcia, E., Yan, M., Guldbrand, S., Kumar, S., Kwakwa, K., Dunsby, C., Neil, M. A. A., French, P. M. W., “Accelerating single molecule localization microscopy through parallel processing on a high‐performance computing cluster,” J. Microscopy 273 148–160 (2019)

19 Kohler TP, Scholz A, Kiachludis D, Hammerschmidt S. “Induction of Central Host Signaling Kinases during Pneumococcal Infection of Human THP-1 Cells” Front Cell Infect Microbiol. 6: 48, Apr 2016;

20 Ma, Hongqiang, Xu, Jianquan, and Liu, Yang. “WindSTORM: Robust Online Image Processing for High-throughput Nanoscopy.” Science Advances 5.4 (2019): Eaaw0683. Web.

21 S. J. Pan and Q. Yang, “A Survey on Transfer Learning,” in IEEE Transactions on Knowledge and Data Engineering, vol. 22, no. 10, pp. 1345–1359, Oct. 2010, 10.1109/TKDE.2009.191

