## Supplementary Information for "Robust optical autofocus system utilizing neural networks trained for extended range and time-course and automated multiwell plate imaging including single molecule localization microscopy"

| | 1 | 2 | 3 | 4 | 5 | 6 | 7 | 8 | 9 | 10 | 11 | 12 |
|---|---|---|---|---|---|---|---|---|---|---|---|---|
| A | 0.0 | -10.7 | -24.02 | -34.75 | -43.73 | -51.49 | -52.44 | -54.4 | -51.29 | -47.55 | -37.5 | -28.61 |
| B | -9.31 | -20.7 | -34.41 | -43.62 | -52.44 | -60.64 | -63.11 | -65.2 | -63.27 | -58.71 | -50.66 | -42.23 |
| C | -12.53 | -25.73 | -38.32 | -49.04 | -57.75 | -65.64 | -69.39 | -72.68 | -71.9 | -69.34 | -63.48 | -57.03 |
| D | -13.28 | -26.99 | -41.3 | -53.35 | -62.77 | -70.1 | -74.33 | -78.63 | -78.71 | -78.34 | -75.02 | -71.37 |
| E | -14.14 | -26.67 | -43.48 | -56.04 | -65.63 | -73.93 | -79.88 | -85.89 | -85.6 | -86.19 | -83.99 | -82.79 |
| F | -15.01 | -27.91 | -44.33 | -58.22 | -69.08 | -79.39 | -87.69 | -90.07 | -94.68 | -93.98 | -94.31 | -96.45 |
| G | -24.5 | -35.3 | -50.81 | -63.52 | -76.7 | -88.97 | -97.78 | -101.25 | -106.33 | -107.91 | -107.88 | -110.77 |
| H | -36.11 | -43.02 | -55.94 | -69.69 | -84.99 | -98.83 | -105.88 | -111.42 | -116.81 | -121.81 | -121.9 | -124.96 |

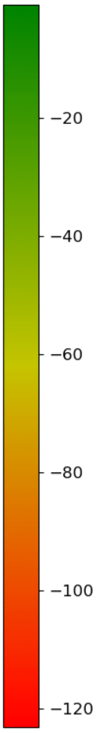


Figure S1. Map of variation of z position of bottom of glass 96-well plate in µm (Brooks life Science Systems, MGB096-1-2-LG-L)

(a) (b)

Figure S2. Videos of exemplar image z-stacks acquired on the autofocus camera as the sample coverslip is translated through the focus of the microscope objective lens: (a) shows a z-stack used to train the "short-range" model over a range of -10µm to +10µm in 50 nm step intervals and (b) shows a z-stack stack used to train the "long-range" model over a range of -100µm to +100µm in 1 µm step intervals.

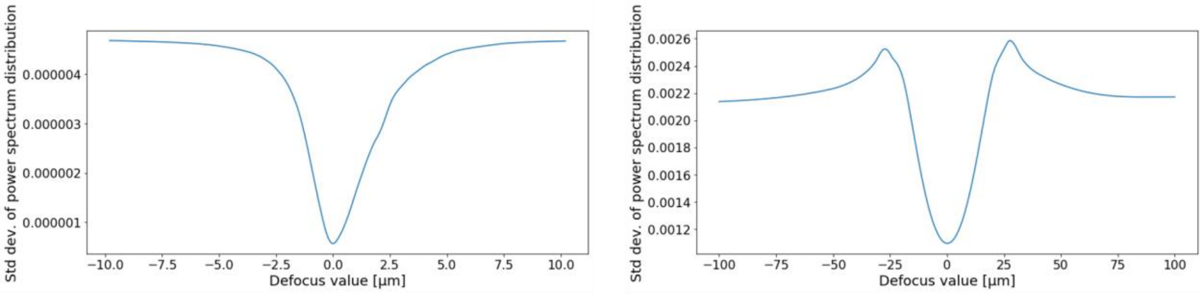


(a) (b)

Figure S3. Exemplar through-focus plots of the standard deviation of the power spectrum of the autofocus camera intensity distribution averaged (a) across the pixel rows for the short-range autofocus data and (b) across the pixel columns for the long-range autofocus data, plotted as a function of z for the images in a z-stack. A Savitzky-Golay filter was passed along these distributions of (averaged) intensities to produce a smoothly varying function.

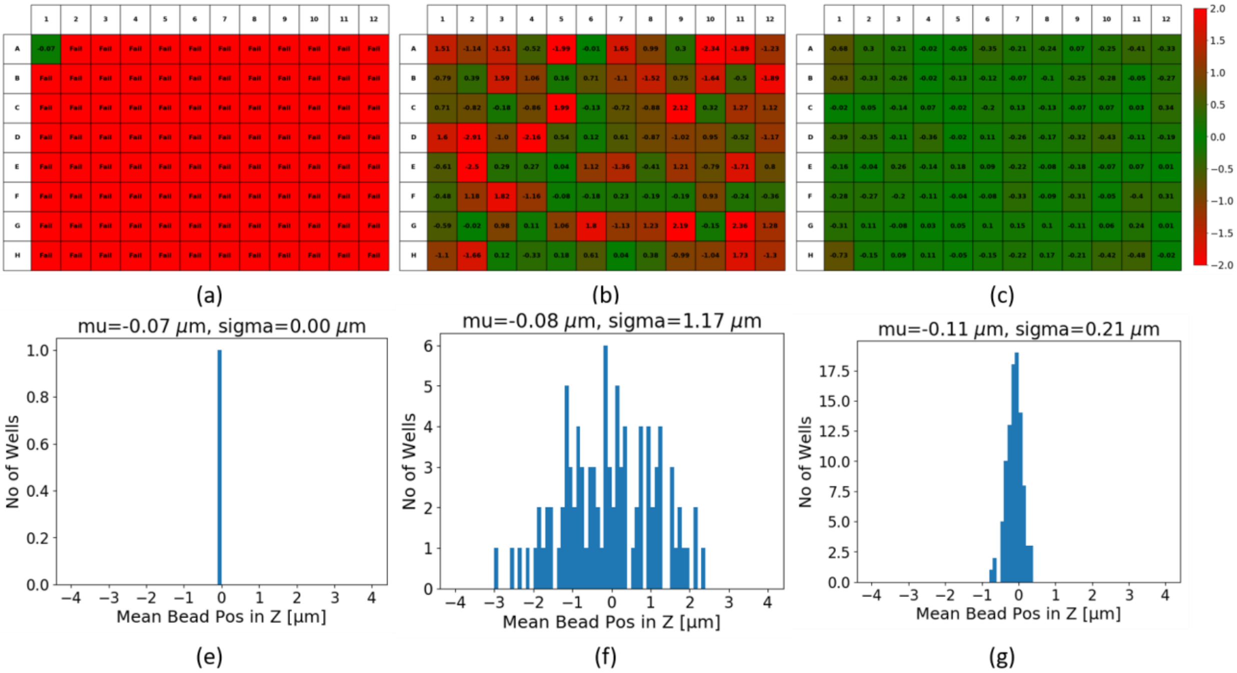


*Figure S4.* Results obtained when automatically imaging 100 nm diameter fluorescent beads (TetraSpeck, T7279) arrayed in a 96-well plate (Brooks life Science Systems, MGB096-1-2-LG-L), for which a through-focus image z-stack was automatically recorded in each well following a horizontal (row-by-row) snake pattern. Figures S3(a-c) show heat maps of the mean defocus of the beads imaged in each well (defocus encoded in colour scale in µm with red indicating that the autofocus failed to locate within +/- 2 µm). The mean defocus of the beads imaged in each cell was calculated using *PSFj*. Where the more precise short-range autofocus training data was used (a, c), the image z-stacks were recorded over a range of -2µm to +2µm in 100 nm steps intervals. For the long-range autofocus (b), the z-stack was recorded between -4µm to +4µm in 100 nm step intervals. In the top row, the heatmaps summarize the results for when (a) only the short-range autofocus, (b) only the long-range autofocus and (c) the combined 2-step autofocus were employed. Where *PSFj* was unable to determine the defocus of the bead images, or where the mean defocus was outside the +/-2 µm range, the heat map presents a red coloured well with either "Fail" or the mean (out-of-range) defocus value. In the bottom row, histograms of the mean defocus for each well, with bin widths of 100 nm, are displayed for the bead image z-stacks for which *PSFj* could determine the defocus. When only using the short-range autofocus (a,e), the system was unable to acquire bead image stacks after the first FOV. When using the long-range autofocus alone (b,f), all the wells were imaged but the standard deviation was 1.17 µm. For the two-step autofocus the standard deviation was 210 nm with bead image stacks being successfully acquired for all of the wells.

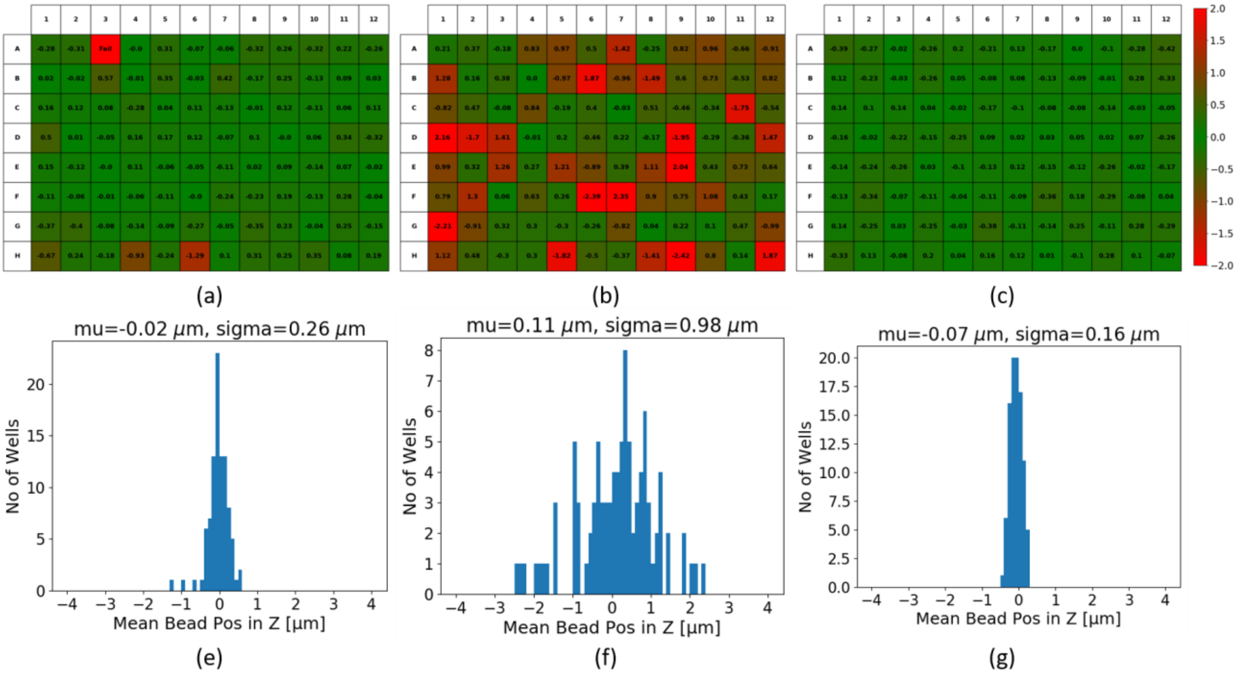


*Figure S5.* Results obtained when automatically imaging 100 nm diameter fluorescent beads (TetraSpeck, T7279) arrayed in a 96-well plate (Brooks life Science Systems, MGB096-1-2-LG-L), for which a through-focus image z-stack was automatically recorded in each well following a vertical (column-by-column) snake pattern. Figures S4(a-c) show heat maps of the mean defocus of the beads imaged in each well (defocus encoded in colour scale in µm with red indicating that the autofocus failed to locate within +/- 2 µm). The mean defocus of the beads imaged in each cell was calculated using *PSFj*. Where the more precise short-range autofocus training data was used (a, c), the image z-stacks were recorded over a range of -2µm to +2µm in 100 nm steps intervals. For the long-range autofocus (b), the z-stack was recorded between -4µm to +4µm in 100 nm step intervals. In the top row, the heatmaps summarize the results for when (a) only the short-range autofocus, (b) only the long-range autofocus and (c) the combined 2-step autofocus were employed. Where *PSFj* was unable to determine the defocus of the bead images, or where the mean defocus was outside the +/-2 µm range, the heat map presents a red coloured well with either “Fail” or the mean (out-of-range) defocus value. In the bottom row, histograms of the mean defocus for each well, with bin widths of 100 nm, are displayed for the bead image z-stacks for which *PSFj* could determine the defocus. When only using the short-range autofocus (e), the system was able to acquire bead image stacks for all but 1 of the FOV’s and the standard deviation of the mean defocus for each well was 264 nm. When using the long-range autofocus alone (f), all the wells were imaged but the standard deviation was 0.982 µm. For the two-step autofocus the standard deviation was 164 nm with bead image stacks being successfully acquired for all of the wells.

**METHODS**

Automated *easySTORM* microscope

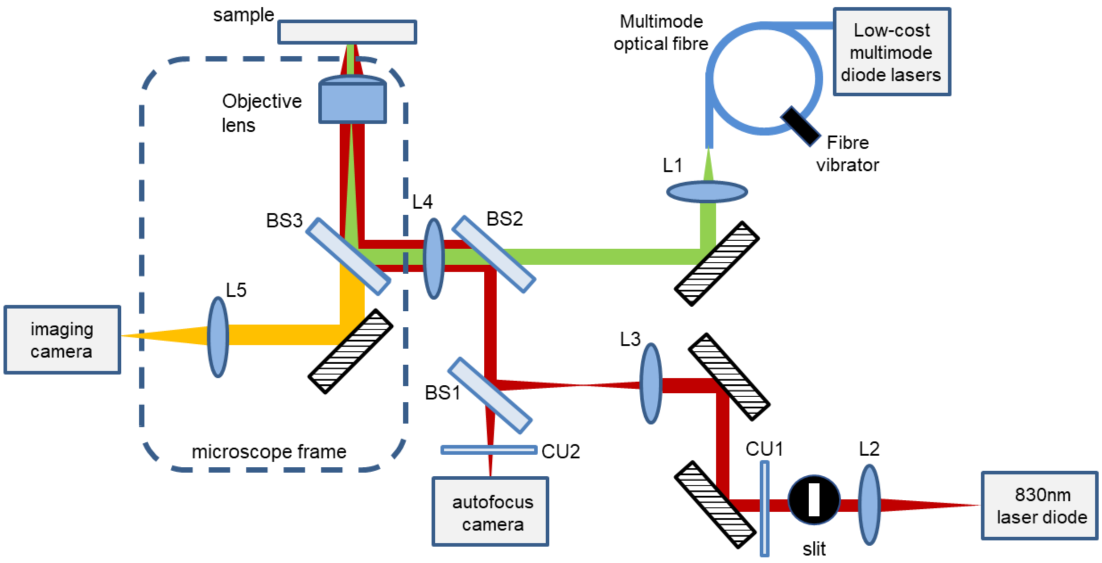


*Figure 1. Schematic of easySTORM microscope utilizing optical autofocus. The objective lens was an oil immersion 1.3 NA 100x lens with a depth of focus of 600 nm. The focal lengths of the other lenses were L1: 50mm, L2: 18.4mm, L3: 100mm, L4: 200mm, L5: 180mm.*

Automated *easySTORM* multiwell plate imaging was implemented as depicted in Figure 1 (shown again for convenience) on a motorised inverted epifluorescence microscope frame (Olympus IX81) with a motorised x-y stage (Marzhauser SCAN IM 112x74, controlled using the Marzhauser TANGO controller). The laser beam for the optical autofocus system was coupled in via the lamp port and directed to the objective lens (Olympus, UplanFl 100x, NA 1.3, oil) via lens L4 (Edmund optics 49-364, 200 nm focal length) and a custom multiline dichroic beamsplitter, BS3. The autofocus laser beam at 830 nm is focused onto the sample and is reflected at the refractive index interface between the glass coverslip and the immersion medium. After recollimation by the objective lens, it is reflected back along its path by the multiline dichroic (BS3). The beam is then focused by and is reflected by the 800 nm short pass beamsplitter (BS2, Edmund Optics 69-220), before being transmitted through the 50/50 beamsplitter, BS1, and focussed onto the autofocus camera (Point Grey, Chameleon3, cm3-U3-31S4M).

The autofocus laser beam was provided by a single spatial mode TO-Can laser diode (HL8338MG), which was able to provide up to a maximum output power of 50mW at 830 nm. The output of this laser was coupled into a single mode optical fibre (FS-SN-4224) using a 10x microscope objective lens and the output of the fibre was collimated using a 18.4 mm aspheric lens (L2, C280TMD-B) such that the diameter of the output collimated beam was 3.68 mm. A Thorlabs adjustable slit (VA100C/M), positioned in the Fourier plane of the aspheric and ideally conjugated to the back focal plane of the objective, was used to produce a quasi-rectangular collimated beam with an aspect ratio of ~1:3 in the x:y axes, corresponding to beam diameters of 1.2:3.68 mm.

A 10 nm bandpass filter at 830 nm, CU1 (Edmund Optics, 65-119) was used to limit the autofocus beam spectrum and the beam was steered using 2 mirrors such that it was aligned along the optical axis of the objective lens after reflecting of BS1 (Edmund Optics, 47-026), and BS2 (Edmund Optics

69-220) and passing through a telescope formed by two achromatic doublet lenses of 100 mm focal length (L3, lens, Thorlabs, AC254-100-B-ML) and 200 mm focal length (L4, Edmund Optics 49-364) respectively, in order to double the diameter of the beam along each axis. Importantly, L4 had an anti-reflection coating from 400-1000nm for use with both the 830 nm autofocus beam and the multiline multimodal laser diode sources used for STORM at 462 nm (1.4W, Lasertack LDM-462-1400-C), 520 nm (1W, Lasertack LDM-520-1000-C) and 638 nm (700mW, Lasertack LDM-638-700-C). A 50:50 beamsplitter (BS1) is used to direct part of the back-reflected autofocus laser beam onto the autofocus camera. The multiline excitation source is designed to provide widefield illumination for STORM using high power low-cost multimode laser diodes coupled into a 105 µm core multimode fibre (M105L02S-A, NA=0.22). The multimode fibre is vibrated to time average speckle the in optical fibre output such that uniform illumination from the fibre is achieved across the whole FOV[1]. The light emitting from the multimode fibre is collimated by an achromatic doublet (L1) of 50 mm focal length (Thorlabs, AC254-050-A-ML), which has an anti-reflection coating between 400-700nm. The collimated excitation beam of ~22 mm diameter is then transmitted through the 800 nm short pass beamsplitter, BS2, and focussed to the back aperture of the objective by the 200 mm focal length achromatic lens (L4) via a reflection at the multiline dichroic beamsplitter (BS3). The focussed spot size of the excitation beam at the back aperture of the objective lens has a diameter of ~420 µm, which results in wide-field uniform illumination at the sample plane over a FOV of ~220 µm diameter. The fluorescence signal from the sample is transmitted through the multiline dichroic (BS3) and emission filter and is relayed by an Olympus tube lens (L5, 180 mm focal length) onto the imaging camera, (Photometrics Iris9 sCMOS).

Cell culture and preparation for imaging

WM266.4 melanoma cells were reverse transfected with OnTArget*Plus* SMARTpools targeting ARHGEF9, ANILLIN, RAC1, CDC42, or RHOA (Dharmacon cat # L-020314-00-0005, cat # L-006838-00-0005, cat # L-003560-00-0005, cat # L-005057-00-005 and cat # L-003860-00-0005) at a concentration of 20 µM in a 6 well plate. Transfections were carried out using Lipofectamine RNAimax (Invitrogen) according to the manufacturer's instructions. 48 hours later cells were trypsinised and resuspended in DMEM, and approximately 5000 cells in 200 µl medium were transferred to a 96 well glass bottom microwell plate (MatriPlate) pre-coated with 10ug/ml fibronectin (Sigma-Aldrich). Ten wells per condition were plated were left to spread for 2.5 hours at 37°C prior to drug treatment. Duplicate wells were treated with 100 µM ARP2/3 inhibitor[2,3] ( CK-666, Sigma-Aldrich), which keeps the complex in an inactive state and inhibits actin nucleation, and SMIFH2[4],[5] (Formin FH2 Domain Inhibitor, Abcam), which prevents formin-mediated actin nucleation (50 mM). Cells treated with CK-666 were left to recover 30 minutes before fixation. Cells were fixed by removing medium and adding 200 ul of pre-warmed 4% methanol free formaldehyde (Thermo) for 10 min at RT. After washing 3X with PBS, cells were permeabilized with 0.1% Triton X-100 (Sigma-Aldrich) for 10 mins at RT. Cells were blocked for 1 hour at RT with 0.5% BSA in PBS. After washing 3X with PBS, primary antibody (Anti-Paxillin, BD Biosciences) was added in 200 µL block solution (1:500), and the plate was sealed and incubated overnight at 4°C. Following 3X washes in PBS, secondary antibody (Alexa Fluor® 488 goat anti mouse IgG, Invitrogen) was added (1:1000) and incubated for 2 hrs at RT. Plate was then washed 2X in PBS, and phalloidin-Atto 647 (Invitrogen) was added (1:5000) in PBS and incubated for 20 minutes at RT. Plate was then washed 2X in PBS and filled with 200 µl PBS before imaging. During imaging, WM266.4 melanoma cells were maintained in Dulbecco's Modified Eagle Medium (DMEM) supplemented with 1% Penicillin + Streptomycin and 10% heat-inactivated fetal bovine serum (HI FBS; GIBCO) cultured at 37°C in 5% $CO_2$.

The THP-1 cells presented in Figure 9 were arrayed in a 96 well plate supplemented with 10% (v/v) fetal calf serum (FCS) (Bioserum), 2mM L-glutamine (Sigma Aldrich) and 10 mg/ml Penicillin/Streptomycin (Sigma Aldrich) at 37oC, 5% (v/v) CO2. Approximately 50,000 cells per well were seeded into each well and were treated with 10 ng/ml phorbol 12-myristate 13-acetate (PMA)

(Sigma Aldrich) for 48h. They were then incubated with heat-killed Streptococcus pneumoniae labelled with AF488 for up to 60 minutes in 15 minute intervals, before washing to remove non-bound bacteria. Cells were then fixed with 100% (v/v) ice-cold methanol for 10min, washed three times with 0.05% (v/v) Triton-X in PBS (permeabilization solution), then blocked for 2h with permeabilization solution supplemented with 10% (v/v) FCS (blocking solution). The cells were then stained with primary monoclonal antibody for alpha-tubulin (Sigma Aldrich) at a 1:1000 dilution in blocking solution overnight at 4°C. They were then washed 3 times with permeabilization solution before incubation with an anti-mouse secondary antibody conjugated with AF647 (ThermoFisher Scientific) at a 1:2000 dilution in blocking solution for 20 minutes at room temperature. Cells were then washed 3 times with permeabilization solution and final washed 3 times with PBS prior to imaging. The imaging STORM buffer was made fresh prior to each imaging session, consisting of 50 mM mercaptamine (Sigma-Aldrich), 10 mM DL-Lactate (Sigma-Aldrich) in PBS. Subsequently, 10 µl of Oxyrase Enzyme (Sigma-Aldrich) was added to each well before being sealed with Parafilm (Sigma-Aldrich).

easySTORM image data analysis

The easySTORM data represented in figure 8 corresponded to fields of view of 125.8 x125.8 µm with typical data volumes of 21.7 Gb per field of view. These data sets were processed using a parallelised implementation of ThunderSTORM run on 4 nodes of an HPC cluster[6]. The ImageJ plug-in for this HPC implementation of ThunderSTORM is available at: https://github.com/ImperialCollegeLondon/HPC_STORM

The easySTORM data represented in figure 9 corresponded to fields of view of 125.8 x125.8 µm with typical data volumes of 10.8 Gb per field of view. These data sets were processed using WindSTORM[7] implemented to run on an HPC cluster.

### References


[1] Kwakwa et al., "easySTORM: a robust, lower-cost approach to localisation and TIRF microscopy", J. Biophotonics 9 (2016) 948–957, http://dx.doi.org/10.1002/jbio.201500324

[2] Hetrick et al. "Small molecules CK-666 and CK-869 inhibit actin-related protein 2/3 complex by blocking an activating conformational change." Chem Biol. 2013 May 23;20(5):701-12. doi: 10.1016/j.chembiol.2013.03.019.

[3] Yamagishi et al. Use of CK-548 and CK-869 as Arp2/3 complex inhibitors directly suppresses microtubule assembly both in vitro and in vivo. Biochem Biophys Res Commun. 2018 Feb 12;496(3):834-839. doi: 10.1016/j.bbrc.2018.01.143.

[4] Rizvi et al. Identification and characterization of a small molecule inhibitor of formin-mediated actin assembly. Chem Biol. 2009 Nov 25;16(11):1158-68. doi: 10.1016/j.chembiol.2009.10.006

[5] Kim, Jo et al. Small Molecule Inhibitor of Formin Homology 2 Domains (SMIFH2) Reveals the Roles of the Formin Family of Proteins in Spindle Assembly and Asymmetric Division in Mouse Oocytes. PLOS ONE (2015) 10(4): e0123438. https://doi.org/10.1371/journal.pone.0123438

[6] Munro, I., Garcia, E., Yan, M., Guldbrand, S., Kumar, S., Kwakwa, K., Dunsby, C., Neil, M. A .A., French, P. M. W., "Accelerating single molecule localization microscopy through parallel processing on a high-performance computing cluster," J. Microscopy 273 148-160 (2019)

[7] Ma, Hongqiang, Xu, Jianquan, and Liu, Yang. "WindSTORM: Robust Online Image Processing for High-throughput Nanoscopy." Science Advances 5.4 (2019): Eaaw0683. Web.
