## Supplementary figures and images for "Robust optical autofocus system utilizing neural networks trained for extended range and time-course and automated multiwell plate imaging including single molecule localization microscopy"

### Supplemental figure 2a

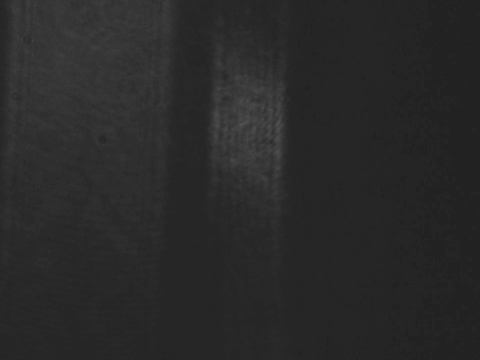

### Supplemental figure 2b

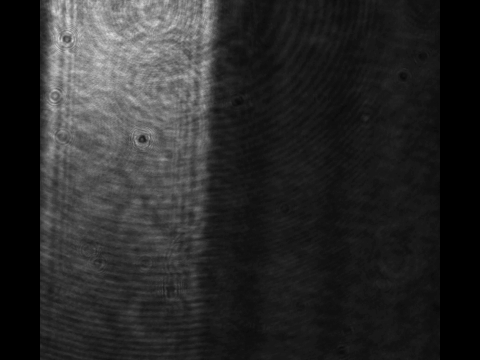
